# Spatial distribution and spread potential of sixteen *Leptospira* serovars in a subtropical region of Brazil

**DOI:** 10.1101/559609

**Authors:** Manuel Jara, Luis E. Escobar, Rogério O. Rodriges, Alba Frias, Juan Sanhueza, Gustavo Machado

## Abstract

Leptospirosis is a bacterial disease that represents a major problem in animal and public health due to its high prevalence and widespread distribution. This zoonotic disease is most prevalent in tropical environments where conditions favor pathogen survival. The ecological preferences of *Leptospira* serovars are poorly understood, limiting our knowledge of where and when outbreaks can occur, which may result in misinformed prevention and control plans. While the disease can occur consistently in time and space in tropical regions, research on the ecology of Leptospirosis remains limited in subtropical regions. This research gap regarding *Leptospira* ecology brings public and veterinary health problems, impacting local economies. To fill this gap of knowledge, we propose to assess geographic and ecological features among *Leptospira* serovars in a subtropical area of Brazil where Leptospirosis is endemic to (i) highlight environmental conditions that facilitate or limit *Leptospira* spread and survival and (ii) reconstruct its geographical distribution. An ecological niche modeling framework was used to characterize and compare *Leptospira* serovars in both geographical and environmental space. Our results show that, despite the geographic overlap exhibited by the different serovars assessed, we found ecological divergence among their occupied ecological niches. Ecological divergences were expressed as ranges of potential distributions and environmental conditions found suitably by serovar, being Sejroe the most asymmetric. Most important predictors for the potential distribution of most serovars were soil pH (31.7%) and landscape temperature (24.2%). Identification of environmental preferences will allow epidemiologists to better infer the presence of a serovar based on the environmental characteristics of regions rather than inferences based solely on historical epidemiological records. Including geographic and ecological ranges of serovars also may help to forecast transmission potential of *Leptospira* in public health and the food animal practice.

## INTRODUCTION

Leptospirosis is a major public health issue due to its high incidence and worldwide distribution (Bharti et al., 2003; Abela-Ridder et al., 2010; Adler and Moctezuma, 2010). Leptospirosis is a zoonotic disease endemic in tropical regions where environmental conditions favor the survival of the bacteria along the year and outside the host (Bharti et al., 2003). Tropical regions often concentrate the highest density of domestic, wild animals and humans (Stevens, 1989; Morand and Poulin, 1998; Gaston, 2000), facilitating interspecies transmission of *Leptospira*, the causative agent of leptospirosis (Adler and Moctezuma, 2010). *Leptospira* serovars have showed to be highly influenced by environmental conditions (Lau et al., 2010; Ivanova et al., 2012). For example, temperature and precipitation (Desvars et al., 2011; Chadsuthi et al., 2012), high humidity and heavy rainfall (Barcellos and Sabroza, 2001; Goarant et al., 2009), runoffs, soil pH, and primary productivity, all have been associated with *Leptospira* occurrence (Smith et al., 1961; Fajriyah et al., 2017; Rahayu et al., 2018).

Approximately 1.03 million cases of leptospirosis are reported globally each year, from which 58,900 are deaths (Costa et al., 2015). Likewise, the global burden estimated in Disability Adjusted Life Years (DALY) per annum for this disease was 2.9 million, showing the great economic impact of leptospirosis worldwide (Torgerson et al., 2015). Leptospirosis is no longer listed among the neglected tropical diseases prioritized by the world health organization (Molyneux et al., 2017). Instead, it is now considered a re-emerging infectious disease linked to a combination of factors, including intensification of livestock production, and limited access to health provision for animals and humans, and environmental change (Mwachui et al., 2015; Hotez, 2016; Goarant et al., 2019). For example, leptospirosis risk is amplified by the frequency of extreme climatic events and major changes in land use (Pappas et al., 2008; Picardeau, 2015). Within these environmental parameters, Brazil is among the top 17 countries in the world with the highest incidence of human leptospirosis (Pappas et al., 2008).

In subtropical regions, the ecology of leptospirosis is generally assumed to be consistent with tropical regions. However, subtropical regions may have considerable environmental differences that may limit effectiveness of control strategies developed for tropical conditions. The Brazilian state of Rio Grande do Sul is located in a subtropical region of southern Brazil. This state has a dense livestock production and one of the highest horse populations in South America (SEAPI-RS., 2018). Additionally, Rio Grande do Sul has the 5/26 highest incidence of human leptospirosis in Brazil (4.7 cases per 10,000 habitants; Ministério da Saúde do Brasil, 2018), representing ∼15% of the total cases (Pacheco and Caldas, 2012). Rio Grande do Sul was also identified among the top five states where improvement on leptospirosis surveillance, control, and elimination must be prioritized in Brazil (Baquero and Machado, 2018). Thus, there is a critical need to identify and anticipate areas and conditions more likely suitable for leptospirosis in this subtropical region.

A comprehensive understanding of the geographic distribution and environmental factors that facilitate *Leptospira* infections will help to inform intervention and prevention strategies for humans and animals (Grooms, 2006; Lilenbaum and Martins, 2014; Sánchez-Montes et al., 2015; Zhao et al., 2016). Such assessments have been widely applied in epidemiology through ecological niche modeling (ENM) in disease ecology. ENM explores geographic and ecological patterns of vectors, hosts or pathogens distribution, and transmission (Peterson, 2006). This approach has shown effectiveness under diverse applications to fundamental ecological questions such as areas at risk of disease infection (Machado et al., 2018), likely pathogen spillover to humans (Peterson, Martínez-Campos, Nakazawa, & Martínez-Meyer, 2005; Samy, Thomas, Wahed, Cohoon, & Peterson, 2016) and environmental factors linked to infectious diseases (Jia and Joyner, 2015; Sallam et al., 2017). Thus, ENM has proven to be a powerful approach to reconstruct the likely factors shaping infectious diseases distributions.

This study aims to identify the geographic and ecological conditions where *Leptospira* serovars occur under subtropical conditions. Using an ENM framework, we characterized and compared the environmental features of *Leptospira* serovars to determine their potential geographic distribution, environmental preferences, and likely hotspots of serovars diversity in the study area. Our approach has the potential to facilitate the development of intelligence-based based leptospirosis surveillance for public health and veterinary epidemiology in this and other subtropical regions.

## MATERIAL AND METHODS

The study design included a modeling framework based on the chosen study area for model calibration, selection, and evaluation, followed by data collection, curation, and standardization (i.e., field work to collect samples, laboratory work for serovar identification, and environmental variables collection and management) (Fig. 1).

**Figure 1.**
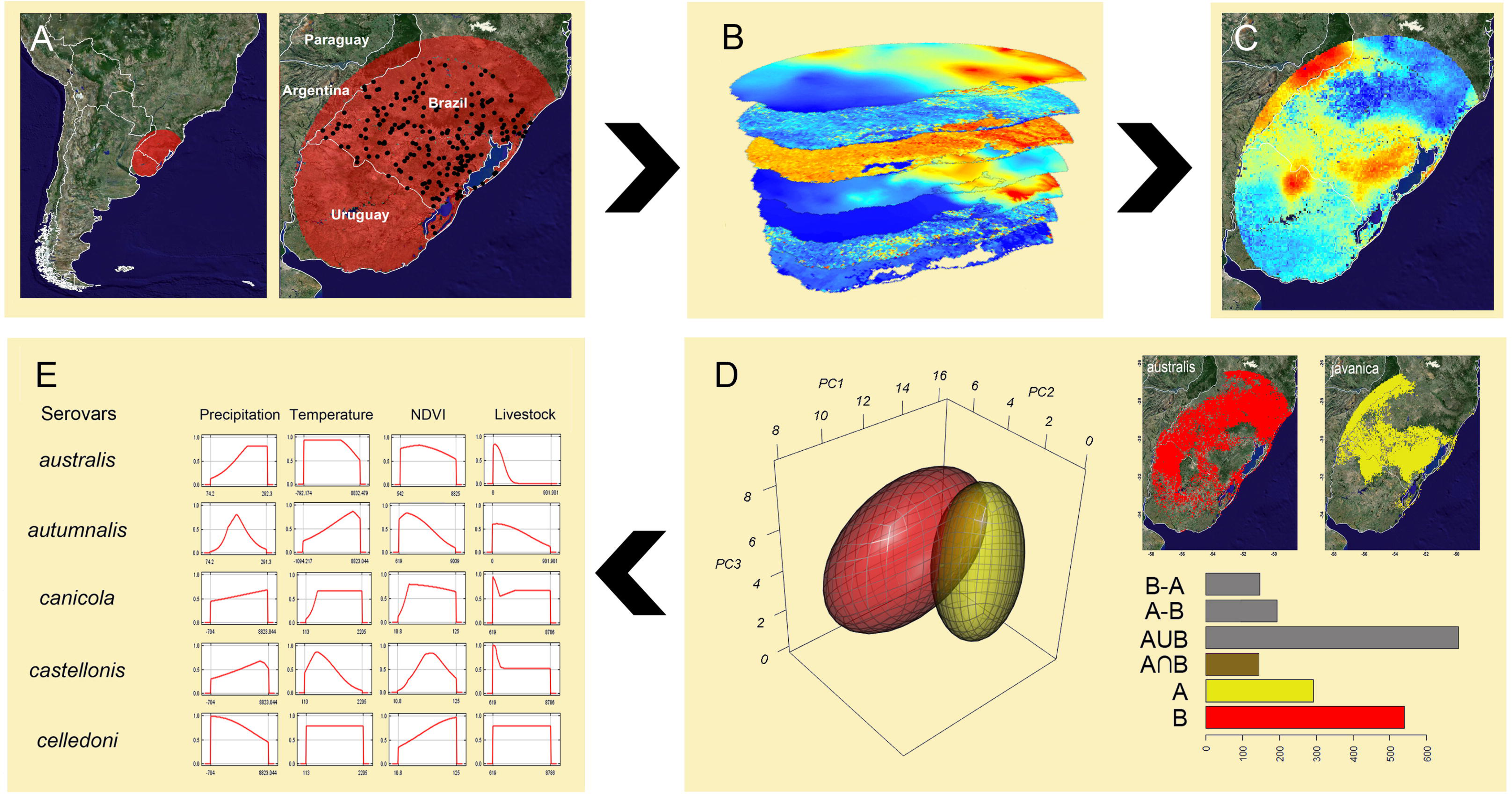
Workflow of the modeling process. **A**) Data collection in the modeling area defined based on biogeographic barriers (**M** region *sensu* (Soberon and Peterson, 2005)), where black dots represent the *Leptospira* occurrence, while the red color shows the study area; **B**) environmental variables relevant for *Leptospira* survival and transmission; **C**) Reduction of number and correlation among environmental variables, model selection, final model calibration; **D**) Assessments of ecological similarity among *Leptospira* serovars, where and the two maps exemplify the potential distribution of two serovars in the geographic space, while 3D plot in the environmental space. Histogram shows ENM comparison between the two serovars based on their environmental space; **E**) Identification of most important environmental for each serovar. NDVI= Normalized Difference Vegetation Index. Red lines represent the response curves that show how each environmental variable affects the ENM prediction for each *Leptospira* serovars, where environmental variables (x axis) and suitability values (y axis) are described for each serovar (left column).

### Data collection

Rio Grande do Sul has ∼103,000 registered horse farms, with a population of more than 550,000 horses (SEAPI-RS., 2018). Our primary dataset comes from a cross-sectional study where 1,010 animals were sampled from 341 farms randomly selected across the state. In each farm, horses were blood sampled to detect previous *Leptospira* exposure. Details of the sampling and laboratory analyses conducted are available elsewhere (Weiblen et al., 2016). Briefly, farms were randomly selected from the total number of farms that had at least one equid older than six months of age (n = 103,170), the number of farms to sample was stratified according to the horse population present in each of the administrative regions of the state (Weiblen et al., 2016). Samples were tested for *Leptospira* antibodies using the microscopic agglutination test (MAT) based on live antigens (Faine et al., 1999; Adler, 2014). Briefly, five 2-fold dilutions of serum samples from 1:25 to 1:4000 were used. Samples were tested for 16 serovars: *Leptospira interrogans* serovar Australis (Australis), *Leptospira interrogans* serovar Autumnalis (Autumnalis), *Leptospira interrogans* serovar Sejroe (Sejroe), *Leptospira interrogans* serovar Canicola (Canicola), *Leptospira interrogans* serovar Ballum (Ballum), *Leptospira interrogans* serovar Celledoni (Celledoni), *Leptospira interrogans* serovar Copenhageni (Copenhageni), *Leptospira borgpetersenii* Javanica (Javanica), *Leptospira interrogans* serovar Grippotyphosa (Grippotyphosa), *Leptospira interrogans* serovar Hardjo (Hardjo), *Leptospira interrogans* serovar Hebdomadis (Hebdomadis), *Leptospira interrogans* serovar Icterohaemorrhagiae (Icterohaemorrhagiae), *Leptospira interrogans* serovar Pomona (Pomona), *Leptospira interrogans* serovar Pyrogenes (Pyrogenes), *Leptospira interrogans* serovar Tarassovi (Tarassovi) and *Leptospira interrogans* serovar Wolffi (Wolffi) (Adler, 2014; Filho et al., 2014; Alves et al., 2016; Dreyfus et al., 2018). The antigens were stored at 28°C from 5 to 10 days in EMJH (Ellinghausen and MCcullough, 1965) culture (Difco^®^-USA) that was enriched with bovine albumin fraction V (Inlab^®^-Brasil) (Ellinghausen and McCullough, 1965). Serum samples were considered positive when MAT titers were ≥ 100. The ultimate reactive serovar was determined by the election of the highest titer that was presented. In the presence of coagglutinations, all serovars that were involved were considered positive (see Fig. S1 to see the spatial information related to positive farms per serovar).

### Selection of the model calibration region

To define the study area extent for model calibration for each *Leptospira* serovar, we followed the framework proposed by (Soberon and Peterson, 2005), which restricts the ENM to ecological features of plausible biological relevance for the organism in question, the resolution of the environmental variables based on the geographic error, and the extent of the region where the organisms are able to disperse based on biogeographic barriers (see **M** in the BAM framework in (Soberon and Peterson, 2005). We assumed a geographic error < 30 m considering that we used GPS devices to estimate the coordinates of each sample and biome regions as biogeographic barriers since they represent homogeneous climatic and landscape composition (Lomolino et al., 2010; Soberón, 2010). This resulted in an **M** in the ecoregion of the Uruguayan savanna (Olson et al., 2001) (see Fig. 1A). This geographic delimitation, **M**, allows to determine the spread potential of the *Leptospira* populations in the study area. Thus, our models are representative of this study area extent.

### Ecological Niche Models (ENMs)

The environmental variables used to estimate the distribution of *Leptospira* were selected based on the described requirements of the bacterium, including survival in specific landscapes with suitable temperature and humidity and presence of livestock (Wint and Robinson, 2007). To reconstruct the landscape structure we used Normalized Difference Vegetation Index (NDVI), a satellite-derived variable resembling vegetation phenology and primary productivity commonly used in ENM (Cook et al., 2008; Fajriyah et al., 2017). We also used annual mean temperature, precipitation, runoff (index that quantity of water discharged in surface streams), and wetness index (defined as a steady-state wetness index), since higher incidences of leptospirosis are related to warmer temperatures (Lau et al., 2010; Desvars et al., 2011; Chadsuthi et al., 2012) and humid environments (Barcellos and Sabroza, 2001; Pappachan et al., 2004; Desvars et al., 2011; Ivanova et al., 2012). In addition, we also included soil pH, since previous studies have explored the importance of this variable in the survival of the bacteria outside the host (Smith et al., 1961; Saito et al., 2013; Schneider et al., 2018). Environmental variables were used at ∼5 km of spatial resolution at the equator (see Table S1 for details). Livestock presence was represented by density of horses, cattle, and pigs (Gilbert et al., 2018), which are known reservoirs and amplifiers of leptospirosis (Lo et al., 2006). To mitigate multicollinearity between environmental variables, we used VIF (Variance Inflation Factors) implemented in the “usdm” R-package (Naimi and Araújo, 2016); excluding highly correlated variables from the model (VIF > 7), since this a signal of strong collinearity (Chatterjee and Hadi, 2015). A detailed description of each environmental variables, such as, description, source, reference and VIF value are presented in (Table S1).

ENMs were developed using a presence-background method that estimates environmental suitability via an index of similarity that resembles a heterogeneous occurrence process or logistic regression function (Phillips et al., 2006; Phillips and Dudík, 2008). Specifically, we used Maxent algorithm with clamping and extrapolation turned off (i.e., no prediction outside the range of environmental conditions used during calibration) (Elith et al., 2010; Anderson, 2013; Owens et al., 2013). To determine the model parametrization with the best fit to the data available, we assessed Maxent models for each serovar under different regularization multiplier values (0.1, 0.3, 0.5, 0.7, 0.9, 1.3, 1.5, 1.7, 1.9 and 2) (Warren and Seifert, 2011). At the same time, we explored all feature combinations ranging from a single feature, linear (L), quadratic (Q), product (P), threshold (T) and hinge (H) (Muscarella et al., 2014), to all feature combinations possible (i.e., 5!). Models were selected based on Akaike Information Criterion (AIC) values, specifically δAICc=0.

To facilitate interpretations of final models, we used the logistic output as a proxy of environmental suitability (Phillips and Dudík, 2008), which we normalized in the final models to suitability ranging from 0 to 100 for easier visualization of values. Additionally, suitable areas for each serovar were estimated as a Boolean (presence/absence) map that was thresholded based on the minimum training presence method to generate binary maps without omission error (Pearson et al., 2006). Serological results of serovars were pooled to general genus-level models (see Table S2 for the best for each set of occurrence data). These models were generated following the protocols described above but focused on the percentage of the variable importance estimated by Maxent. To determine the hotspots of *Leptospira* serovars richness, we ensembled all binary models by using Spatial Analysis in Macroecology (SAM) software (Rangel et al., 2010), (available at https://ecoevol.ufg.br/sam).

In addition, we showed how each environmental variable affects the Maxent ENM prediction for each serovar, representing how the predicted probability of presence changes as each environmental variable is varied.

### Ecological niche similarity among serovars

We assessed ecological similarities among models of *Leptospira* serovars by using four methods based on geographic and environmental dimensions. First, similarity was measured using the Schoener’s *D* index (Schoener, 1968) that measures similarity between two ENMs in geographic space based on probabilities outputs being similar in terms of the environmental conditions available to them (Rödder and Engler, 2011). Schoener’s *D* was estimated by comparing Maxent-generated ENMs against a null distribution of default Maxent models, resulting in similarity values ranging from 0, non-similar, to 1, highly similar. We followed the protocols described by (Warren et al., 2008) and (Warren et al., 2010). Second, we used the Jaccard similarity index (Jaccard, 1912) that assesses similarity between two ENMs in environmental space by measuring the volume and overlap of two ENMs (Escobar et al., 2015). Volume of environmental space occupied by each serovar was estimated in three forms to capture variability among estimates. First, volume of ENMs was estimated in NicheA software (Qiao et al., 2016), available at http://nichea.sourceforge.net/function_niche_overlap.html. Briefly, original environmental variables were collapsed into three environmental dimensions to reduce redundancy and dimensionality. Then, volume was measured based on the environments occupied by each *Leptospira* serovar in terms of a minimum-volume ellipsoid and a convex-polyhedron. Finally, volume was estimated for all serovars combinations using “hypervolume” package in the R software (Blonder et al., 2014). This method relies on a Gaussian kernel density estimation procedure based on the Silverman method (Silverman, 1986), measuring the geometry of the multidimensional hypervolume from the original variables standardized (Blonder et al., 2014). The Jaccard similarity index based on NicheA and hypervolume values provides an accurate measure of the geometrical relationships between serovars distribution in a multidimensional space (Goodall, 1966; Real and Vargas, 1996). In summary, we generated one similarity estimation in geographic space (Schoener’s *D*) and three estimations in environmental space (Jaccard indices from the minimum-volume ellipsoid, convex-polyhedron, and Gaussian kernel density).

## RESULTS

### Spatial patterns of Leptospira serovars distribution

Approximately 45% of the total *Leptospira* serovars circulating in the study area were Hebdomadis, Tarassovi, Pyrogenes. (Fig. S1). Occurrence of the 16 *Leptospira* serovars showed considerable asymmetries among their geographic distribution (Fig. S1).

### Ecological niche models (ENMs)

ENM results showed that central-northern areas in this study had suitable conditions for Autumnalis, Canicola, Copenhageni, Hardjo, Icterohaemorrhagiae, and Wolfii. Contrarily, central-western areas were suitable for Calledoni, Grippotyphosa, Javanica, Hebdomadis, and Pyrogenes were most common. Australis, Pomona and Tarassovi were found to prefer the northern area, while Serjroe, and Castellonis concentrated its potential distribution in the western and eastern regions, respectively (Fig 2).

**Figure 2.**
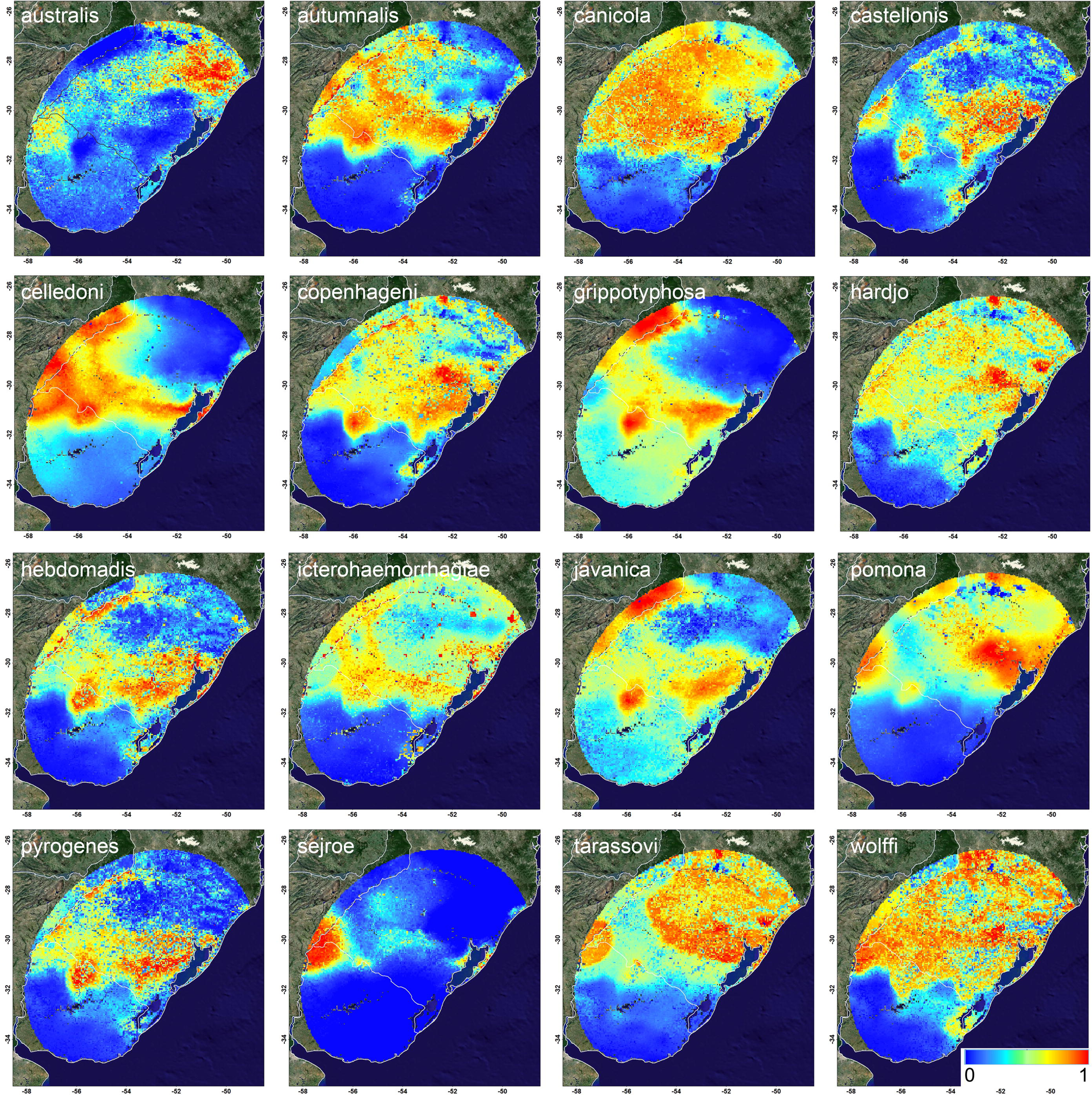
Ecological niche model (ENM) predictions of *Leptospira* serovars in Southern Brazil. Warmer colors show areas with higher probability of presence. Background layer represents the earth in true color based on NASA’s Terra satellite image for better visualization. Source https://neo.sci.gsfc.nasa.gov/.

These distributional differences among serovars were also reflected in the geographic patterns observed in the hotspot areas per serovar. Similarly, the model ensemble comprised specific areas of agreement of *Leptospira* suitability. For example, the region bordering northern Argentina and areas between Caxias do Sul and Taquari River, were hotspot for *Leptospira* likely exposure. Additional areas of *Leptospira* exposure-risk were found in Taquarombo, Uruguay, and Lagoa Mirim, between Brazil and northeastern Uruguay (Fig. 3A). The visualization of the ensembled model also in a multidimensional environmental space, revealed that *Leptospira* occurred under consistent and trackable environmental conditions, however, available conditions were more diverse and broader than those occupied by the pathogen (Fig. 3B).

**Figure 3.**
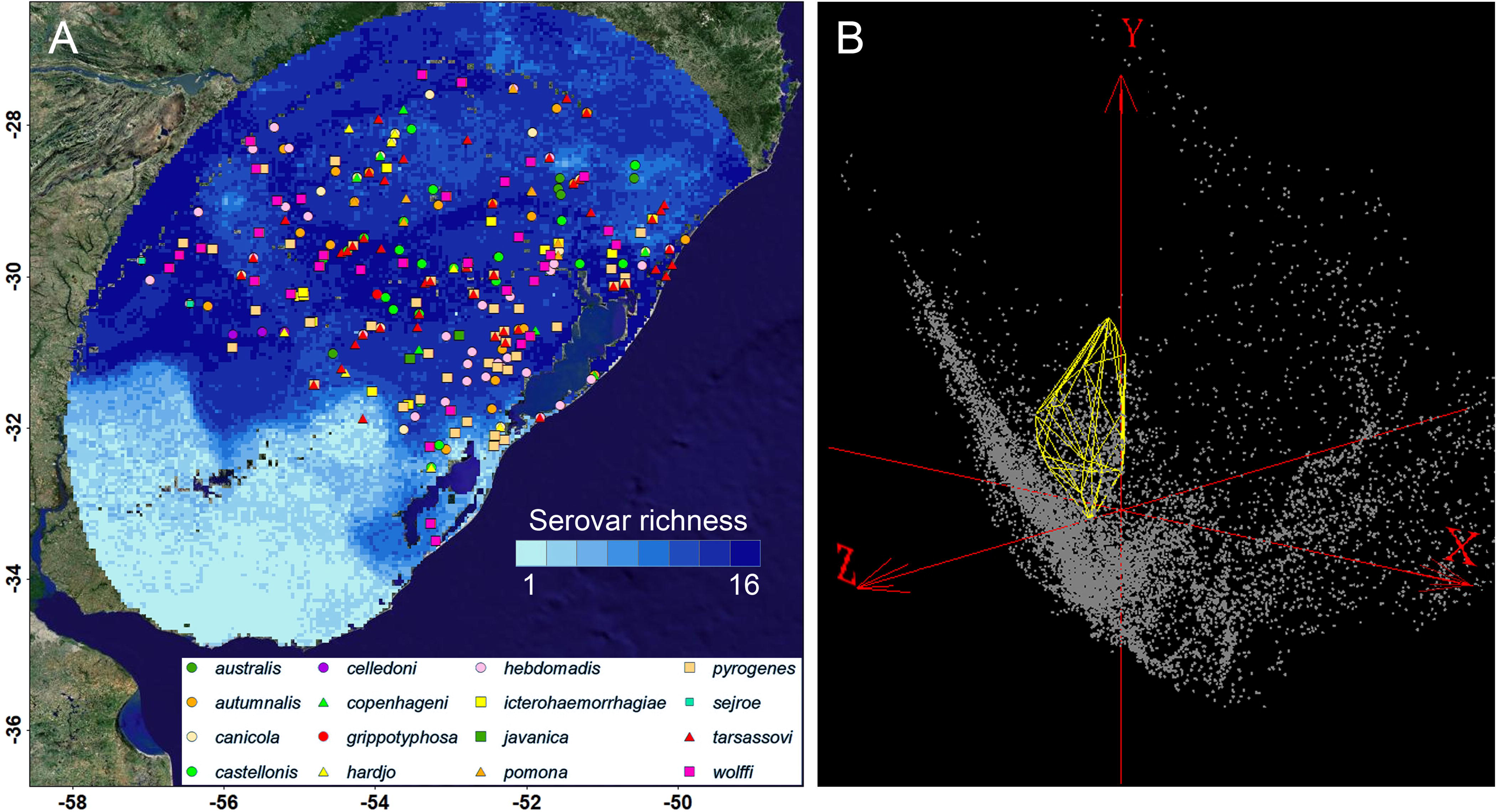
Model ensembled of *Leptospira* serovars in a subtropical region. **A**) Model ensemble in geographic space based on binary models summed to identify areas of highly (dark blue) and low (light blue) agreement among models. Dark areas denote areas found consistently suitable for *Leptospira* and therefore, for plausible exposure infection. **B**) Model ensemble in environmental space based on binary models summed to identify the environmental conditions occupied by the sero-positive cases (yellow convex polyhedron). Grey dots represent the environmental available conditions in **M**, axes are the first principal components from the original environmental variables (see Table S1).

### Environmental drivers of serovars potential distribution

The final *Leptospira* ENM showed that soil pH (31.7%) and mean annual temperature (24.2%) were the most influential predictors associated with *Leptospira* sero-positivity (Table 1). Response curves also suggested that as pH and temperature increased linearly the suitability index for *Leptospira* presence (Fig. 4). We identified the importance of variables related with humidity wetness; index; runoff and precipitation in the distribution of Australis, Autumnalis, Canicola, Celledoni, Pomona, Wolffi, Sejroe, and Tarassovi. Likewise, NDVI was the main predictor for Grippotyphosa sero-positivity, while livestock production was observed to be the most important predictor for serovar Sejroe sero-positivity (Table 1).

**Table 1.** Variable importance for all *Leptospira* and for each serovar. Warmer colors represent higher levels of importance (%).

| Serovar | Temperature | Wetness index | Soil pH | NDVI | Runoff | Precipitation | Livestock |
| --- | --- | --- | --- | --- | --- | --- | --- |
| *Leptospira* (all) | 24.2 | 9.2 | 31.7 | 4.4 | 6.2 | 12.0 | 12.3 |
| Australis | 22.1 | 2.8 | 19.9 | 8.2 | 42.4 | 0.5 | 4.1 |
| Autumnalis | 3.6 | 20.8 | 35.9 | 7.4 | 10.7 | 11.5 | 10.1 |
| Canicola | 0.0 | 27.7 | 55.4 | 2.5 | 0.0 | 0.3 | 14.1 |
| Castellonis | 7.8 | 12.0 | 56.7 | 4.7 | 6.3 | 9.2 | 3.3 |
| Celledoni | 0.0 | 0.0 | 54.7 | 0.0 | 20.2 | 25.1 | 0.0 |
| Copenhageni | 5.7 | 0.0 | 66.5 | 0.0 | 0.0 | 6.7 | 21.1 |
| Grippotyphosa | 14.7 | 0.0 | 25.9 | 59.4 | 0.0 | 0.0 | 0.0 |
| Hardjo | 32.8 | 13.7 | 39.8 | 0.0 | 0.0 | 0.0 | 13.7 |
| Hebdomadis | 10.4 | 13.9 | 63.1 | 0.7 | 11.2 | 0.0 | 0.7 |
| Icterohaemorrhagiae | 0.0 | 2.6 | 72.2 | 4.4 | 0.0 | 0.9 | 19.9 |
| Javanica | 12.3 | 18.1 | 51.2 | 5.1 | 0.0 | 10.6 | 2.7 |
| Pomona | 0.0 | 24.3 | 44.1 | 0.7 | 0.0 | 30.9 | 0.0 |
| Pyrogenes | 34.7 | 14.5 | 42.6 | 0.0 | 1.6 | 3.1 | 3.5 |
| Sejroe | 0.1 | 0.1 | 0.2 | 0.1 | 8.9 | 44.8 | 45.9 |
| Tarassovi | 10.3 | 0.1 | 20.0 | 5.5 | 59.9 | 2.2 | 2.1 |
| Wolffi | 21.4 | 26.9 | 40.9 | 2.3 | 0.0 | 6.6 | 1.9 |

**Figure 4.**
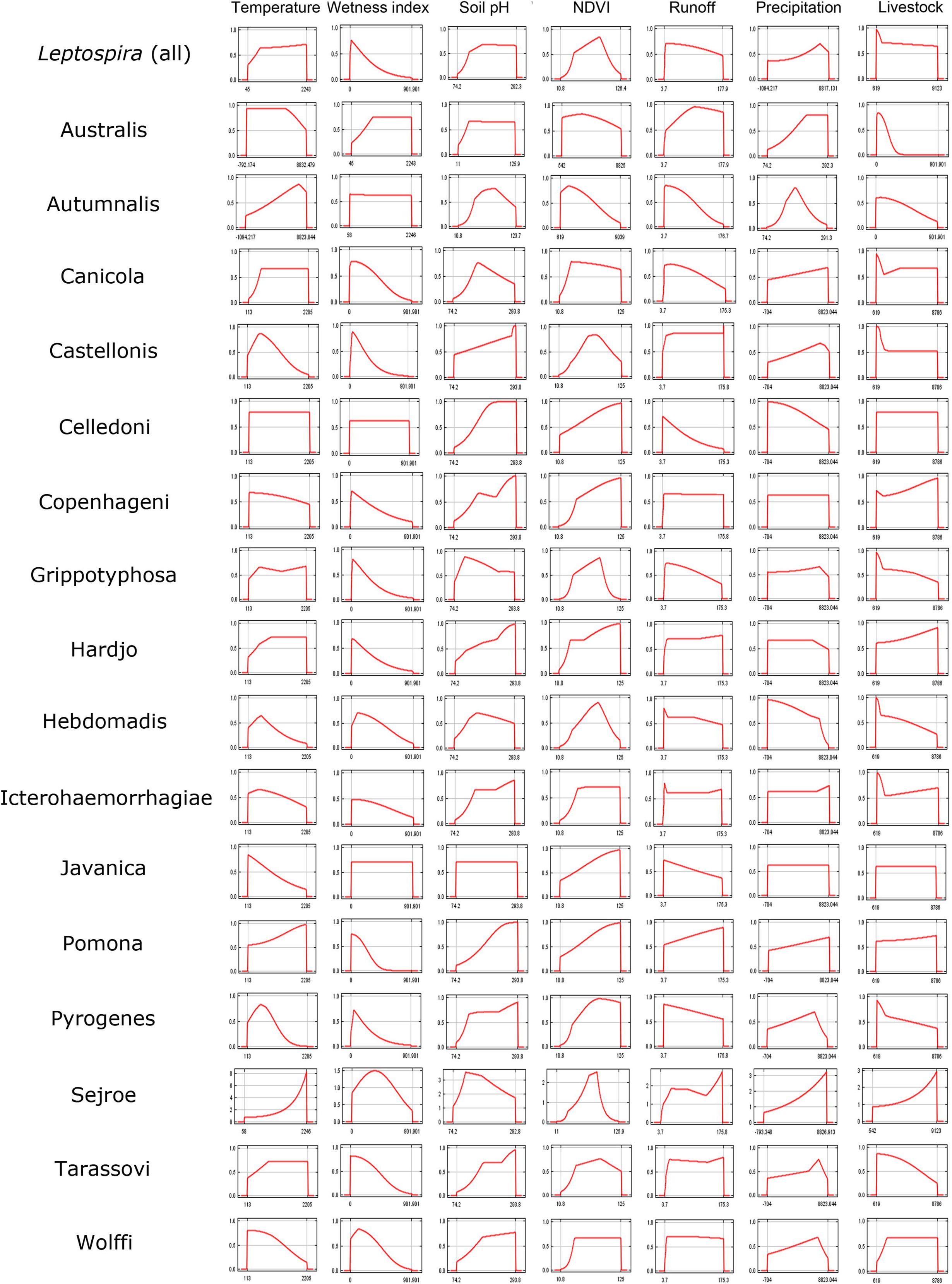
Response curves of the different environmental variables by *Leptospira* serovars. Response curves (red line) estimated based on Maxent ENM predictions. Environmental variables (x axis) and suitability values (y axis) are described for each serovar (left column). Units, source, and details of each variable are found in Table 1.

### Ecological similarities among Leptospira serovars

The observed values of Schoener’s (*D*) similarity tests showed niche similarity of all serovars, which tend to overlap on average 0.68 ± 0.2, ranging from 0.09 (Sejroe and Castellonis) to 0.97 (Icterohaemorrhagiae and Javanica). In all cases, the serovar that showed the most asymmetric results was Sejroe, where the highest observed similarity values were under 0.15 (Fig. 5A and Table S3). These considerable variations in the ecological niches between serovars were also observed in the NicheA results, showing that the ecological niche of *Leptospira* is characterized by asymmetries in the distribution of the different serovars in the environmental space. The niche overlap was on average 0.16 ± 0.08 “Convex polyhedron” (CP) (Fig. 5B, Table S4) and 0.16 ± 0.09 “Minimum Volume Ellipsoid” (MVE) (Fig. 5C, Table S5). These values ranged from 0.01 (CP= Sejroe and Pomona, MVE= Sejroe and Grippotyphosa) to 0.3 (CP= Copenhageni and Tarassovi) and 0.38 (MVE= Tarassovi and Wolffi). Similarly to what was previously observed, large asymmetries were observed by the “hypervolume” approach. Serovars tend to be similar on average 0.11 ± 0.09, with values ranging from 0 (Grippotyphosa with Copenhageni, Hardjo, Icterohaemorrhagiae, Javanica, Pomona and Sejroe) to 0.26 (Copenhageni and Pomona) (Fig. 5D and Table S6).

**Figure 5.**
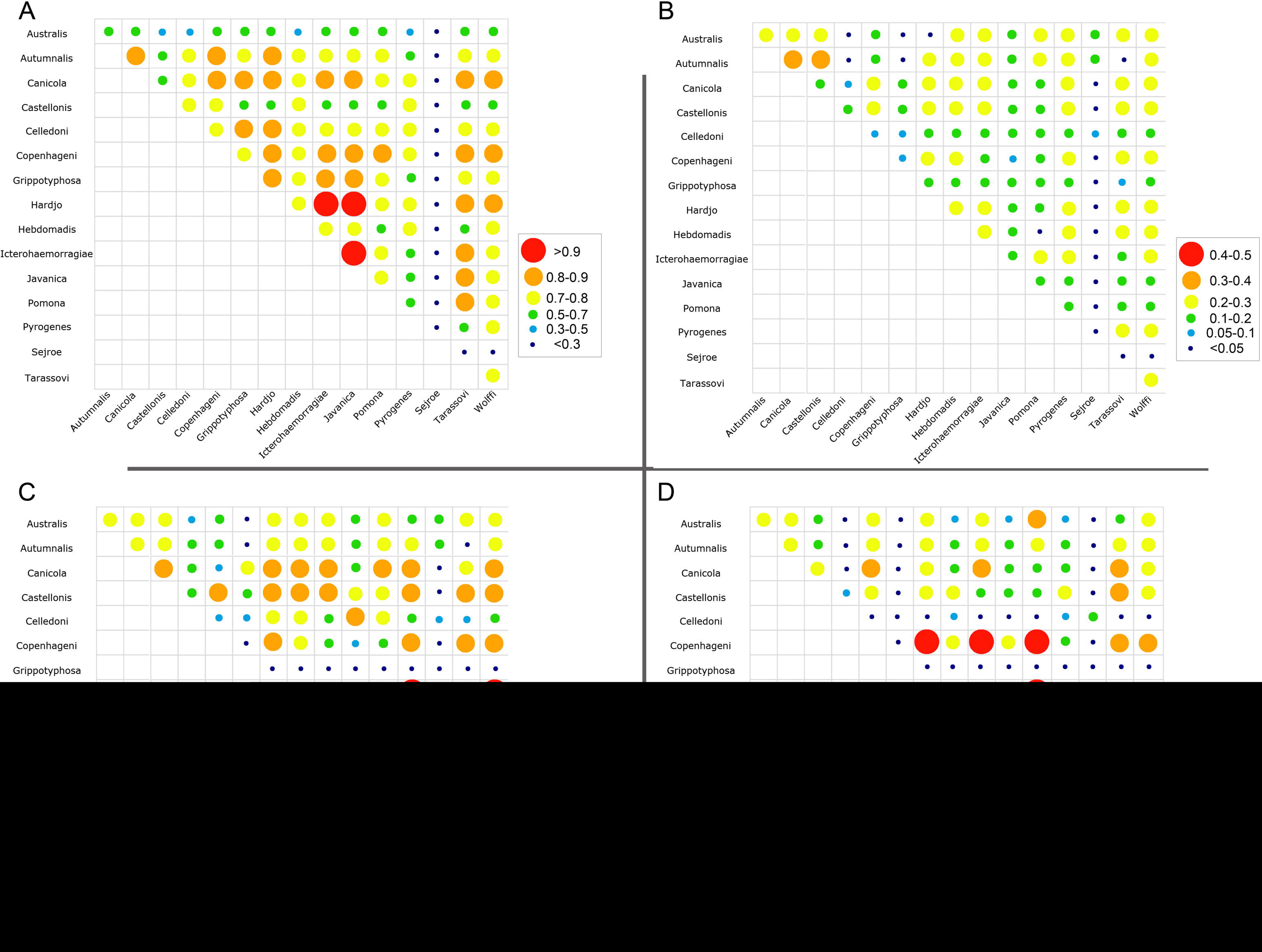
Ecological niche similarity between *Leptospira* serovars based on four model similarity metrics. **A**) Schoener’s *D* index, **B**) convex polyhedron, **C**) Minimum Volume Ellipsoid and **D**) hypervolume.

Overall, higher asymmetries in the ecological niche were evidenced between the majority of *Leptospira* serovars, these results were highly contrasted with the results based on the geographical space (Schoener’s (*D*) index) (Fig. 5A and Table S3), while, the lowest similarity values were observed in the comparison based on convex polyhedron (Fig. 4B).

## DISCUSSION

This study combines geographic and ecological approaches to characterize eco-epidemiology patterns of *Leptospira* sero-positivity in horses at serovar level. This cross-sectional study allowed for the identification of geographic and ecological preferences of *Leptospira* serovars. Despite the geographic similarities exhibited by each serovar, they showed different environmental preferences, evidencing the diversity of environmental conditions where *Leptospira* exposure can occur. Recent efforts have been made to understand the environmental tolerances of *Leptospira* at serovar level (Fouts et al., 2016; Guernier et al., 2017; Jaeger et al., 2018; Zarantonelli et al., 2018), as well as to identify potential risk areas for future leptospirosis outbreaks (Sánchez-Montes et al., 2015; Zhao et al., 2016). However, there have been no studies able to forecast regions where high-transmission risk exists and where disease surveillance and control strategies (e.g., vaccination) would have better impact. Our multidimensional approach (i.e., geographic and environmental dimensions) represents an important stepping-stone in the study and understanding of *Leptospira* ecology not only for identifying risk areas for different serovars but also for the development of new strategies to understand the ecological drivers of *Leptospira* presence.

Our results showed considerable differences in the ecological landscape features of the distribution of each *Leptospira* serovar. This could be explained by the fact that different *Leptospira* lineages can survive and adapt to different environmental conditions. Historically, leptospirosis has been widely associated to warm and humid conditions (Barcellos and Sabroza, 2001; Trueba et al., 2004; Ivanova et al., 2012; Saito et al., 2013; Schneider et al., 2018). Our results support these previous findings since the variables related to temperature and precipitation were considered highly important predictors for the potential distribution of most of the serovars, especially for Australis, Hardjo, Pyrogenes and Tarassovi. Looking into detail, our results showed a positive relationship between precipitation and a higher probability of presence of *Leptospira*. However, we also found negative relationship of *Leptospira* presence with soil humidity and runoff, which could be explained by the type of soil in the area: bentonite clay, which is an amplifier of this pathogen since it absorbs half of *Leptospira* in suspension (Smith et al., 1961). On the other hand, responses curves of each serovar associated to temperature did not show a clear, consistent pattern.

Recently, Schneider et al., (2018), highlighted the importance of pH in the survival of this bacteria in soil, which was strongly supported by our findings. We found that soil pH was the main predictor for 12 of the 16 serovars examined. Response curves evidenced a positive relationship between soil pH and the probability of presence of most of *Leptospira* serovars, except for Autumnalis, Canicola, Hebdomadis and Javanica.

Niche similarity tests based on environmental space (convex polyhedron, MVE, and hypervolume) revealed high asymmetries between the majorities of *Leptospira* serovars. These results were highly contrasted with what we observed in the geographical space (Schoener’s *D* index), where most of the serovars tends to overlap their distributions. The niche similarity tests offer biological realism to the different models by giving access to a broader perspective that support the idea of phylogenetic niche conservatism among the *Leptospira* lineages studied (Escobar, Qiao, Phelps, Wagner, & Larkin, 2016; Martinez-Meyer, Diaz-Porras, Peterson, & Yanez-Arenas, 2012; Yañez-Arenas, Peterson, Mokondoko, Rojas-Soto, & Martínez-Meyer, 2014). The importance and significance of the use of these similarity tests at serovar level relies in the fact that disease transmission is the product of complex interactions that involves ecological, evolutionary, and epidemiological processes (Fountain-Jones et al., 2018; Galvani, 2003; Peterson, 2006).

Risk of horizontal gene transmission can occur between serovars (Ren et al., 2003; Haake et al., 2004), which can facilitate shifts in virulence (Dzidic and Bedekovic’, 2003; Khairani-Bejo et al., 2004; Salyers et al., 2004; Adler, 2014). Thus, the possibility of multiple serovars cohabiting in the same location increases the possibility of gene transfer making our serovar richness maps informative to design *Leptospira* monitoring plans if areas of higher disease-emergence risk (Fig. 3).

ENM is used to characterize environmental requirements of species and their potential distribution s (Peterson, 2014; Peterson & Vieglais, 2001; Qiao et al., 2018). These analyses have been applied for a wide variety of epidemiological purposes such as the prediction of species invasions into novel areas (Benedict et al., 2007; Machado et al., 2018), anticipation of disease emergence (Peterson, Bauer, & Mills, 2004; Williams & Peterson, 2009), and forecast of the impact of climate change on future emerging disease distributions (González et al., 2010; Gálvez et al., 2011; Daszak et al., 2013; De Oliveira et al., 2017; Baquero and Machado, 2018). Our approach represents a novel application of ENM aimed to generate new knowledge about the ecology of *Leptospira* at serovar level. However, more efforts are necessary to determine if our findings are consistent in different biogeographic regions (e.g., tropical, temperate). Finally, we faced limitations through the development of this study, mainly related to the species sampled. More specifically, to obtain the most accurate representation of *Leptospira* circulation in the landscape there would be necessary to assess the presence of *Leptospira* serovars in wildlife and the environment to provide and integrative estimation of the geographic and environmental risk (Albert et al., 2009).

## CONCLUSION

In this study, we identified the geographic and environmental signatures of *Leptospira* serovars in a subtropical region in southern Brazil. We determined the geographic and ecological characteristics influencing the current and potential distributions of all *Leptospira* serovars tested providing new ecological and epidemiological knowledge about *Leptospira* lineages circulation in animal populations. We found specific environmental preferences of serovars, most serovars were limited by soil pH and mean annual temperature. The maps generated in this study also denote the local and regional hotspots of disease transmission risk, useful to design evidence-based disease prevention strategies for effective surveillance and vaccination.

## ACKNOWLEDGEMENTS

We acknowledge the Rio Grande do Sul Official Veterinary Service (SEAPI) and their assistants during the sampling in the field. Special thanks to the official veterinarians from the State of Rio Grande do Sul for their important role in leading the sampling: G. N. Diehl, and L. L. C. dos Santos and L. G. Corbellini from Universidade Federal do Rio Grande do Sul.

## Supporting information

### Supplementary Tables

**Table S1.** Variable selection and Variance Inflation Factor analysis (VIF) to assess spatial multicollinearity.

**Table S2.** Model evaluation results based on Akaike’s information criteria (δAICc = 0). Considering a variety of feature classes (linear= L, product= P, quadratic= Q, threshold= T and hinge= H) and regularization multiplier. AUC=Area Under Curve (Variance).

**Table S3.** Ecological niche similarity between *Leptospira* serovars (based on Schoener’s D index) for *Leptospira* serovars, where Aus = australis, Aut = autumnalis, Can = canicola, Cas = castellonis, Cel = celledoni, Cop = copenhageni, Grip = grippotyphosa, Har = hardjo, Heb = hebdomadis, Ict = icterohaemorrhagiae, Jav = javanica, Pom = pomona, Pyr = pyrogenes, Ser = sejroe, Tar = tarassovi and Wol = wolffi. Warmer colors represent higher similarity values.

**Table S4.** Ecological niche similarity between *Leptospira* serovars (based on Hypervolume approach) for *Leptospira* serovars, where Aus = australis, Aut = autumnalis, Can = canicola, Cas = castellonis, Cel = celledoni, Cop = copenhageni, Grip = grippotyphosa, Har = hardjo, Heb = hebdomadis, Ict = icterohaemorrhagiae, Jav = javanica, Pom = pomona, Pyr = pyrogenes, Ser = sejroe, Tar = tarassovi and Wol = wolffi. Warmer colors represent higher similarity values.

**Table S5.** Environmental niche comparison matrix (Jaccard index) based on minimum volume ellipsoid for *Leptospira serovars*, where Aus = australis, Aut = autumnalis, Can = canicola, Cas = castellonis, Cel = celledoni, Cop = copenhageni, Grip = grippotyphosa, Har = hardjo, Heb = hebdomadis, Ict = icterohaemorrhagiae, Jav = javanica, Pom = pomona, Pyr = pyrogenes, Ser = sejroe, Tar = tarassovi and Wol = wolffi. Warmer colors represent higher similarity values.

**Table S6**. Environmental niche comparison matrix (Jaccard index) based on convex polyhedron for Leptospira serovars, where Aus = australis, Aut = autumnalis, Can = canicola, Cas = castellonis, Cel = celledoni, Cop = copenhageni, Grip = grippotyphosa, Har = hardjo, Heb = hebdomadis, Ict = icterohaemorrhagiae, Jav = javanica, Pom = pomona, Pyr = pyrogenes, Ser = sejroe, Tar = tarassovi and Wol = wolffi. Warmer colors represent higher similarity values.

### Supplementary figures

**Figure S1.** Geographic occurrences of *Leptospira* serovars in Southern Brazil.

**Figure S2.** Binary ENM predictions of *Leptospira* serovars in Southern Brazil.

