## Supplementary material for "Spatial distribution and spread potential of sixteen *Leptospira* serovars in a subtropical region of Brazil"

**Supplementary Tables**

**Table S1.** Variable selection and Variance Inflation Factor analysis (VIF) to assess spatial multicollinearity.

| Variable | Description (unit) | Source | Reference | VIF |
| --- | --- | --- | --- | --- |
| Livestock production | Number of individuals per pixel (5 min of arc) | [https://dataverse.harvard.edu/dataverse/glw_3](http://www.fao.org/ag/againfo/resources/en/glw/GLW_dens.html) | [(Gilbert et al., 2018)](http://www.fao.org/ag/againfo/resources/en/glw/GLW_dens.html) | 5.02 |
| Mean temperature | **°**C | MODIStsp R package and https://neo.sci.gsfc.nasa.gov/ | (Busetto and Ranghetti, 2016) | 2.03 |
| NDVI |  | MODIStsp R package and https://neo.sci.gsfc.nasa.gov/ | (Busetto and Ranghetti, 2016) | 1.10 |
| Mean precipitation | mm | <https://climate.northwestknowledge.net/TERRACLIMATE/index_directDownloads.php> | [(Abatzoglou et al., 2018)](https://climate.northwestknowledge.net/TERRACLIMATE/index_directDownloads.php) | 3.63 |
| Runoff | mm | <https://climate.northwestknowledge.net/TERRACLIMATE/index_directDownloads.php> | [(Abatzoglou et al., 2018)](https://climate.northwestknowledge.net/TERRACLIMATE/index_directDownloads.php) | 2.61 |
| Wetness index | mm | [http://worldgrids.org](http://worldgrids.org/) | [(Hengl et al., 2015)](http://worldgrids.org/) | 1.26 |
| Soil PH |  | [https://www.soilgrids.org/index.html#!/?layer=TAXNWRB_250m&vector=1](https://www.soilgrids.org/index.html" \l "!/?layer=TAXNWRB_250m&vector=1) | Food and Agriculture Organization of the United Nations (FAO) International Soil Reference and Information Centre (ISRIC) | 4.56 |

**Table S2.** Model evaluation results based on Akaike’s information criteria (ΔAICc = 0). Considering a variety of feature classes (linear= L, product= P, quadratic= Q, threshold= T and hinge= H) and regularization multiplier. AUC=Area Under Curve (Variance).

| Serovar | Feature | Regularization Multiplier |
| --- | --- | --- |
| *Leptospira spp.* | PT | 2.0 |
| australis | QTHL | 1.3 |
| autumnalis | LQT | 1.0 |
| canicola | T | 2.0 |
| castellonis | LQ | 0.1 |
| celledoni | T | 1.5 |
| copenhageni | T | 2.0 |
| grippotyphosa | T | 1.7 |
| hardjo | T | 1.5 |
| hebdomadis | LT | 2.0 |
| icterohaemorrhagiae | T | 2.0 |
| javanica | PT | 2.0 |
| pomona | L | 1.0 |
| pyrogenes | T | 1.5 |
| serjroe | L | 0.1 |
| tarassovi | QTHL | 1.7 |
| wolffi | T | 1.0 |

**Table S3.** Ecological niche similarity between *Leptospira* serovars (based on Schoener’s D index) for *Leptospira* serovars, where Aus= australis, Aut= autumnalis, Can= canicola, Cas= castellonis, Cel= celledoni, Cop= copenhageni, Grip= grippotyphosa, Har= hardjo, Heb= hebdomadis, Ict= icterohaemorrhagiae, Jav= javanica, Pom= pomona, Pyr= pyrogenes, Ser= sejroe, Tar= tarassovi and Wol= wolffi. Warmer colors represent higher similarity values.

| Serovar | Aut | Can | Cas | Cel | Cop | Grip | Har | Heb | Ict | Jav | Pom | Pyr | Ser | Tar | Wol |
| --- | --- | --- | --- | --- | --- | --- | --- | --- | --- | --- | --- | --- | --- | --- | --- |
| Aus | 0.62 | 0.63 | 0.44 | 0.48 | 0.55 | 0.65 | 0.67 | 0.44 | 0.51 | 0.56 | 0.59 | 0.44 | 0.10 | 0.66 | 0.59 |
| Aut | * | 0.81 | 0.64 | 0.70 | 0.81 | 0.72 | 0.81 | 0.73 | 0.73 | 0.75 | 0.76 | 0.65 | 0.11 | 0.77 | 0.78 |
| Can | * | * | 0.69 | 0.78 | 0.89 | 0.81 | 0.89 | 0.76 | 0.85 | 0.84 | 0.79 | 0.73 | 0.11 | 0.83 | 0.82 |
| Cas | * | * | * | 0.71 | 0.70 | 0.66 | 0.65 | 0.71 | 0.63 | 0.63 | 0.69 | 0.73 | 0.09 | 0.68 | 0.67 |
| Cel | * | * | * | * | 0.79 | 0.84 | 0.81 | 0.77 | 0.78 | 0.78 | 0.76 | 0.73 | 0.11 | 0.79 | 0.77 |
| Cop | * | * | * | * | * | 0.78 | 0.86 | 0.75 | 0.82 | 0.80 | 0.82 | 0.75 | 0.12 | 0.84 | 0.85 |
| Grip | * | * | * | * | * | * | 0.86 | 0.72 | 0.87 | 0.87 | 0.75 | 0.69 | 0.09 | 0.78 | 0.73 |
| Har | * | * | * | * | * | * | * | 0.73 | 0.94 | 0.93 | 0.78 | 0.71 | 0.11 | 0.84 | 0.81 |
| Heb | * | * | * | * | * | * | * | * | 0.71 | 0.70 | 0.66 | 0.78 | 0.12 | 0.69 | 0.74 |
| Ict | * | * | * | * | * | * | * | * | * | 0.97 | 0.75 | 0.69 | 0.10 | 0.81 | 0.78 |
| Jav | * | * | * | * | * | * | * | * | * | * | 0.75 | 0.68 | 0.10 | 0.81 | 0.77 |
| Pom | * | * | * | * | * | * | * | * | * | * | * | 0.65 | 0.11 | 0.86 | 0.73 |
| Pyr | * | * | * | * | * | * | * | * | * | * | * | * | 0.09 | 0.68 | 0.74 |
| Ser | * | * | * | * | * | * | * | * | * | * | * | * | * | 0.13 | 0.12 |
| Tar | * | * | * | * | * | * | * | * | * | * | * | * | * | * | 0.79 |

**Table S4**. Environmental niche comparison matrix (Jaccard index) based on convex polyhedron for *Leptospira* serovars, where Aus= *australis*, Aut= *autumnalis*, Can= canicola, Cas= castellonis, Cel= celledoni, Cop= copenhageni, Grip= grippotyphosa, Har= hardjo, Heb= hebdomadis, Ict= icterohaemorrhagiae, Jav= javanica, Pom= pomona, Pyr= pyrogenes, Ser= sejroe, Tar= tarassovi and Wol= wolffi. Warmer colors represent higher similarity values.

| Serovar | Aut | Can | Cas | Cel | Cop | Grip | Har | Heb | Ict | Jav | Pom | Pyr | Ser | Tar | Wol |
| --- | --- | --- | --- | --- | --- | --- | --- | --- | --- | --- | --- | --- | --- | --- | --- |
| Aus | 0.27 | 0.28 | 0.29 | 0.04 | 0.19 | 0.02 | 0.01 | 0.29 | 0.25 | 0.11 | 0.23 | 0.23 | 0.12 | 0.25 | 0.26 |
| Aut | * | 0.30 | 0.32 | 0.05 | 0.17 | 0.03 | 0.28 | 0.28 | 0.26 | 0.14 | 0.22 | 0.27 | 0.12 | 0.05 | 0.26 |
| Can | * | * | 0.20 | 0.09 | 0.20 | 0.14 | 0.29 | 0.21 | 0.26 | 0.13 | 0.19 | 0.26 | 0.01 | 0.22 | 0.26 |
| Cas | * | * | * | 0.17 | 0.21 | 0.12 | 0.20 | 0.27 | 0.22 | 0.16 | 0.19 | 0.25 | 0.02 | 0.23 | 0.23 |
| Cel | * | * | * | * | 0.09 | 0.08 | 0.13 | 0.16 | 0.13 | 0.15 | 0.11 | 0.14 | 0.05 | 0.10 | 0.13 |
| Cop | * | * | * | * | * | 0.07 | 0.21 | 0.21 | 0.17 | 0.09 | 0.12 | 0.23 | 0.01 | 0.30 | 0.20 |
| Grip | * | * | * | * | * | * | 0.13 | 0.10 | 0.18 | 0.13 | 0.19 | 0.12 | 0.00 | 0.09 | 0.12 |
| Har | * | * | * | * | * | * | * | 0.24 | 0.25 | 0.12 | 0.18 | 0.28 | 0.01 | 0.23 | 0.29 |
| Heb | * | * | * | * | * | * | * | * | 0.22 | 0.16 | 0.01 | 0.28 | 0.01 | 0.22 | 0.27 |
| Ict | * | * | * | * | * | * | * | * | * | 0.16 | 0.24 | 0.24 | 0.01 | 0.20 | 0.25 |
| Jav | * | * | * | * | * | * | * | * | * | * | 0.16 | 0.15 | 0.02 | 0.10 | 0.15 |
| Pom | * | * | * | * | * | * | * | * | * | * | * | 0.18 | 0.00 | 0.14 | 0.18 |
| Pyr | * | * | * | * | * | * | * | * | * | * | * | * | 0.01 | 0.24 | 0.29 |
| Ser | * | * | * | * | * | * | * | * | * | * | * | * | * | 0.01 | 0.01 |
| Tar | * | * | * | * | * | * | * | * | * | * | * | * | * | * | 0.21 |

**Table S5.** Environmental niche comparison matrix (Jaccard index) based on minimum volume ellipsoid for *Leptospira serovars*, where Aus= australis, Aut= autumnalis, Can= canicola, Cas= castellonis, Cel= celledoni, Cop= copenhageni, Grip= grippotyphosa, Har= hardjo, Heb= hebdomadis, Ict= icterohaemorrhagiae, Jav= javanica, Pom= pomona, Pyr= pyrogenes, Ser= sejroe, Tar= tarassovi and Wol= wolffi. Warmer colors represent higher similarity values.

| Serovar | Aut | Can | Cas | Cel | Cop | Grip | Har | Heb | Ict | Jav | Pom | Pyr | Ser | Tar | Wol |
| --- | --- | --- | --- | --- | --- | --- | --- | --- | --- | --- | --- | --- | --- | --- | --- |
| Aus | 0.25 | 0.27 | 0.23 | 0.08 | 0.15 | 0.03 | 0.25 | 0.30 | 0.26 | 0.15 | 0.22 | 0.20 | 0.13 | 0.22 | 0.26 |
| Aut | * | 0.28 | 0.24 | 0.10 | 0.12 | 0.04 | 0.23 | 0.24 | 0.25 | 0.18 | 0.22 | 0.22 | 0.15 | 0.04 | 0.23 |
| Can | * | * | 0.21 | 0.12 | 0.06 | 0.19 | 0.26 | 0.22 | 0.27 | 0.15 | 0.23 | 0.25 | 0.02 | 0.19 | 0.25 |
| Cas | * | * | * | 0.11 | 0.21 | 0.12 | 0.25 | 0.30 | 0.21 | 0.16 | 0.20 | 0.27 | 0.03 | 0.21 | 0.26 |
| Cel | * | * | * | * | 0.08 | 0.10 | 0.15 | 0.18 | 0.13 | 0.22 | 0.16 | 0.14 | 0.10 | 0.09 | 0.14 |
| Cop | * | * | * | * | * | 0.03 | 0.20 | 0.19 | 0.14 | 0.07 | 0.10 | 0.21 | 0.01 | 0.28 | 0.21 |
| Grip | * | * | * | * | * | * | 0.05 | 0.04 | 0.05 | 0.02 | 0.04 | 0.03 | 0.01 | 0.04 | 0.05 |
| Har | * | * | * | * | * | * | * | 0.26 | 0.23 | 0.15 | 0.21 | 0.30 | 0.03 | 0.20 | 0.31 |
| Heb | * | * | * | * | * | * | * | * | 0.21 | 0.17 | 0.21 | 0.27 | 0.03 | 0.19 | 0.27 |
| Ict | * | * | * | * | * | * | * | * | * | 0.15 | 0.28 | 0.23 | 0.02 | 0.18 | 0.22 |
| Jav | * | * | * | * | * | * | * | * | * | * | 0.17 | 0.15 | 0.07 | 0.08 | 0.15 |
| Pom | * | * | * | * | * | * | * | * | * | * | * | 0.20 | 0.03 | 0.14 | 0.19 |
| Pyr | * | * | * | * | * | * | * | * | * | * | * | * | 0.02 | 0.21 | 0.31 |
| Ser | * | * | * | * | * | * | * | * | * | * | * | * | * | 0.01 | 0.25 |
| Tar | * | * | * | * | * | * | * | * | * | * | * | * | * | * | 0.38 |

**Table S6. Ecological niche similarity between *Leptospira* serovars (based on Hypervolume approach) for *Leptospira* serovars, where Aus= australis, Aut= autumnalis, Can= canicola, Cas= castellonis, Cel= celledoni, Cop= copenhageni, Grip= grippotyphosa, Har= hardjo, Heb= hebdomadis, Ict= icterohaemorrhagiae, Jav= javanica, Pom= pomona, Pyr= pyrogenes, Ser= sejroe, Tar= tarassovi and Wol= wolffi. Warmer colors represent higher similarity values.**

| Serovars | Aut | Can | Cas | Cel | Cop | Grip | Har | Heb | Ict | Jav | Pom | Pyr | Ser | Tar | Wol |
| --- | --- | --- | --- | --- | --- | --- | --- | --- | --- | --- | --- | --- | --- | --- | --- |
| Aus | 0.16 | 0.17 | 0.11 | 0.04 | 0.16 | 0.02 | 0.18 | 0.08 | 0.15 | 0.08 | 0.21 | 0.08 | 0.02 | 0.10 | 0.15 |
| Aut | * | 0.18 | 0.14 | 0.04 | 0.18 | 0.01 | 0.17 | 0.14 | 0.18 | 0.11 | 0.17 | 0.12 | 0.02 | 0.16 | 0.18 |
| Can | * | * | 0.16 | 0.04 | 0.20 | 0.03 | 0.19 | 0.14 | 0.22 | 0.15 | 0.10 | 0.10 | 0.04 | 0.23 | 0.19 |
| Cas | * | * | * | 0.05 | 0.19 | 0.42 | 0.18 | 0.20 | 0.12 | 0.15 | 0.11 | 0.17 | 0.07 | 0.25 | 0.16 |
| Cel | * | * | * | * | 0.04 | 0.02 | 0.03 | 0.06 | 0.02 | 0.03 | 0.02 | 0.06 | 0.10 | 0.03 | 0.04 |
| Cop | * | * | * | * | * | 0.00 | 0.26 | 0.19 | 0.25 | 0.17 | 0.27 | 0.14 | 0.04 | 0.23 | 0.21 |
| Grip | * | * | * | * | * | * | 0.00 | 0.00 | 0.00 | 0.00 | 0.00 | 0.01 | 0.00 | 0.05 | 0.04 |
| Har | * | * | * | * | * | * | * | 0.16 | 0.21 | 0.13 | 0.26 | 0.13 | 0.04 | 0.21 | 0.22 |
| Heb | * | * | * | * | * | * | * | * | 0.15 | 0.18 | 0.15 | 0.21 | 0.03 | 0.23 | 0.20 |
| Ict | * | * | * | * | * | * | * | * | * | 0.12 | 0.13 | 0.11 | 0.02 | 0.23 | 0.16 |
| Jav | * | * | * | * | * | * | * | * | * | * | 0.07 | 0.16 | 0.07 | 0.18 | 0.15 |
| Pom | * | * | * | * | * | * | * | * | * | * | * | 0.10 | 0.00 | 0.26 | 0.17 |
| Pyr | * | * | * | * | * | * | * | * | * | * | * | * | 0.06 | 0.18 | 0.15 |
| Ser | * | * | * | * | * | * | * | * | * | * | * | * | * | 0.04 | 0.03 |
| Tar | * | * | * | * | * | * | * | * | * | * | * | * | * | * | 0.25 |

**Supplementary figures**

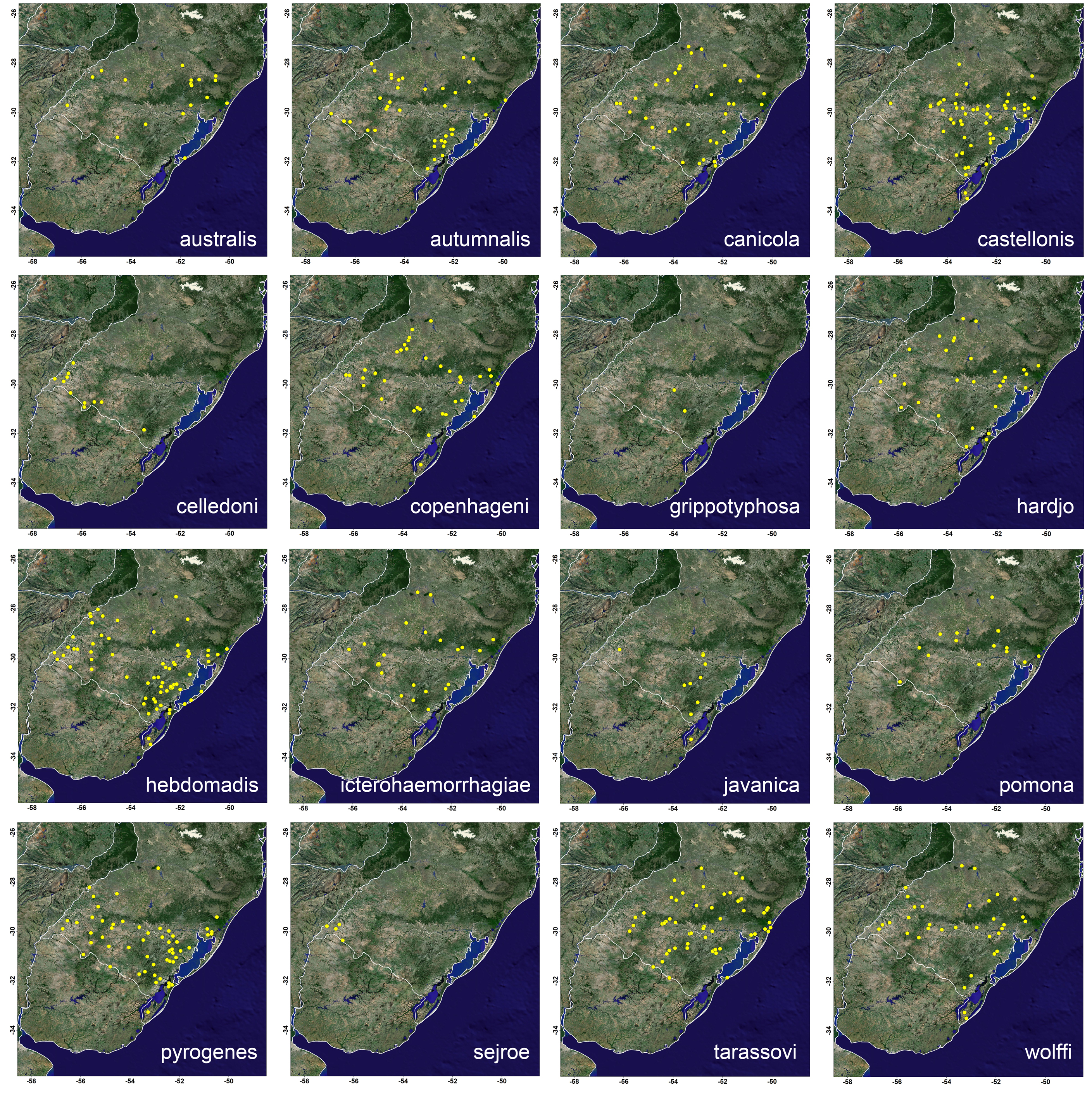

**Figure S1.** Geographic occurrences of Leptospira serovars in Southern Brazil. Background layer represents the earth in true color based on NASA's Terra satellite image, available at: <https://neo.sci.gsfc.nasa.gov/>.

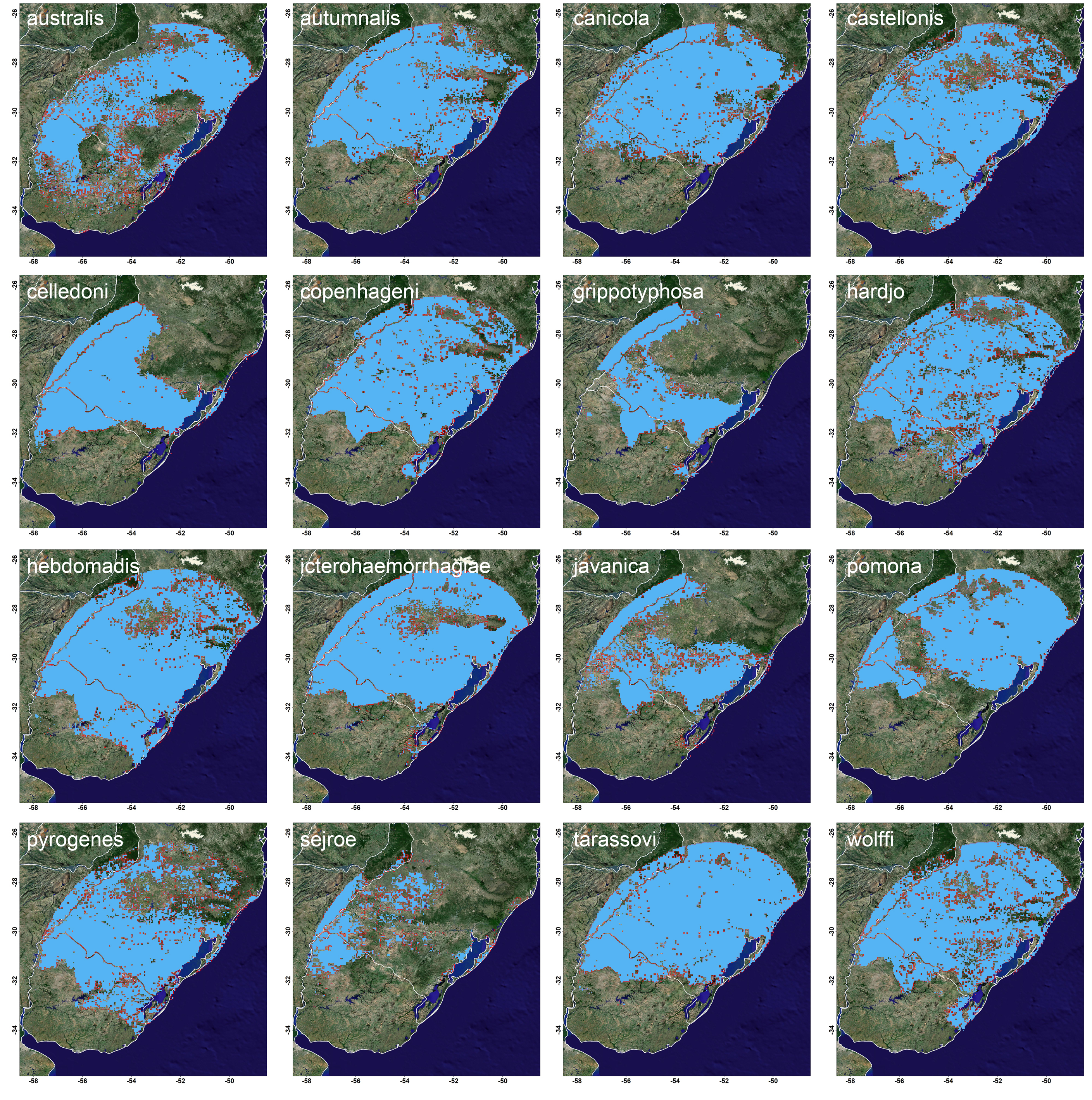

**Figure S2.** Binary ENM predictions of Leptospira serovars in Southern Brazil. Background layer represents the earth in true color based on NASA's Terra satellite image, available at: <https://neo.sci.gsfc.nasa.gov/>.
